# HBIcloud: An Integrative Multi-Omics Analysis Platform

**DOI:** 10.1101/2024.08.31.607334

**Authors:** Shuang He, Yunqing Luo, Wei Dong, Wenquan Wang, Fei Chen

## Abstract

As biological datasets have grown exponentially in size and complexity, there has been an increasing need for integrative tools that can handle diverse data types and facilitate comprehensive analyses. Traditional methods often require significant computational expertise, creating barriers for many researchers. HBIcloud is a comprehensive online platform designed to facilitate multi-omics data analysis by integrating a wide array of tools across genomics, transcriptomics, proteomics, metabolomics, phenomics, and multi-omics integration. Developed to address the growing complexity and volume of biological data, HBIcloud provides researchers with a powerful and user-friendly resource for conducting sophisticated analyses without the need for extensive programming skills. With a total of 94 tools, the platform offers standardized workflows, extensive parameter options, and rich documentation, catering to the diverse needs of the scientific community. The research behind HBIcloud aimed to create a centralized, user-friendly platform that simplifies the analytical process, enabling researchers to focus on scientific discovery rather than technical challenges. By integrating a wide array of tools and offering extensive support and documentation, HBIcloud addresses the critical need for standardized, reproducible workflows in multi-omics research. This paper presents a detailed overview of HBIcloud, highlighting its development background, key features, and its significant contribution to advancing multi-omics research. Furthermore, we discuss the future prospects of HBIcloud, including planned enhancements and its potential for high citation impact within the scientific community. By providing a robust and versatile platform, HBIcloud aims to accelerate discovery and innovation in the field of multi-omics, fostering collaborative research and expanding the boundaries of biological understanding. The official website of HBIcloud is https://bioinformatics.hainanu.edu.cn/HBIcloud/.

## Introduction

Omics technologies have revolutionized biological research^1^, enabling the high-throughput collection of data at multiple biological levels. Genomics^2^, the study of an organism’s complete set of DNA, has provided insights into genetic variations and evolutionary relationships. Transcriptomics^3^, which focuses on RNA transcripts produced by the genome, has revealed gene expression patterns and regulatory mechanisms. Proteomics^4^, the large-scale study of proteins, has elucidated protein functions and interactions. Metabolomics^5^, the comprehensive analysis of metabolites, has highlighted metabolic pathways and their dynamics. Finally, phenomics^6^, the study of phenotypes on a large scale, has linked genotypic variations to observable traits.

Despite the progress in each omics field, a single-layer analysis often falls short of capturing the complexity of biological systems. Integrating these diverse datasets, known as multi-omics integration, is essential for a holistic understanding of biological processes. For instance, integrating genomics and transcriptomics data can identify how genetic variations influence gene expression^7^. Combining proteomics and metabolomics data can elucidate the biochemical pathways regulated by proteins and metabolites^8^. Such integrative approaches can reveal intricate biological networks and uncover novel biomarkers or therapeutic targets.

However, multi-omics integration is fraught with challenges^9, 10^. The heterogeneous nature of omics data, with varying data types, scales, and dimensions, complicates the integration process. Differences in data quality, batch effects, and missing values further add to the complexity. Additionally, the computational and statistical methods required for effective multi-omics integration are often sophisticated^11^, demanding substantial expertise and resources. These challenges create a significant barrier for researchers, especially those lacking advanced bioinformatics skills or access to high-performance computing infrastructure.

Recognizing these challenges, the initial motivation for developing HBIcloud was to democratize access to multi-omics data analysis tools. We aimed to create a platform that not only integrates diverse omics data but also provides user-friendly interfaces and comprehensive support to empower researchers from various backgrounds (Figure 1). HBIcloud was designed to streamline the entire analysis process, from data preprocessing and normalization to integration and visualization, ensuring that high-quality, reproducible analyses are accessible to all researchers.

**Figure 1.**
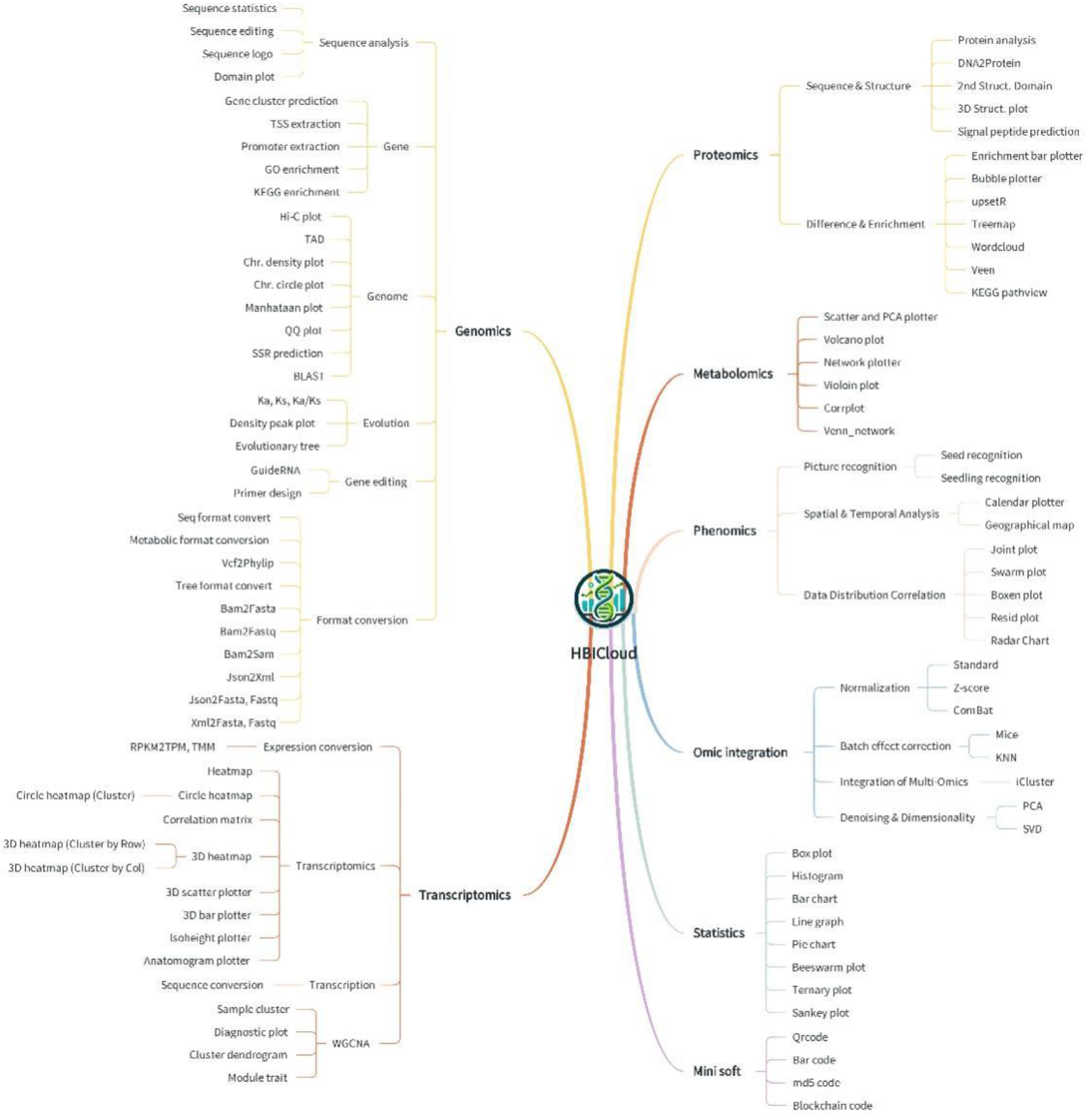
Overview of the 94 tools integrated in HBIcloud. All these tools are classified according to the different omics, including genomics, transcriptomics, proteomics, metabolomics, phenomics, omic integration, as well as statistics and mini soft.

## Materials and methods

### The implementation of HBIcloud

Java (version 17.0.10), jQuery (version 3.2.1), Python (version 3.8) and R (version 4.3.1) serve as the primary technologies for developing the platform. The front-end interface is predominantly crafted using the Thymeleaf framework (version 3.0.12, https://thymeleaf.org/index.html) and Bootstrap (version 5.3.0, https://getbootstrap.com). These tools facilitate data submission by researchers and customization of result presentation. SpringBoot (version 2.4.4, https://docs.spring.io/spring-boot/index.html) functions as the backend web framework, managing the data and parameters submitted by researchers. The analytical outcomes are then relayed to the front-end for researchers to review, with the option to download result files. In the realm of data visualization, HBIcloud leverages ggplot2 (version 3.5.1), matplotlib (version 3.7.5), and seaborn (version 0.13.2). By harnessing the analytical and visualization capabilities of these libraries, researchers can delve into data exploration and analysis in an intuitive manner.

Collectively, the amalgamation of SpringBoot, Thymeleaf, jQuery, Bootstrap, ggplot2, matplotlib, and seaborn empowers HBIcloud to seamlessly amalgamate backend functionalities, front-end development, and dynamic data analysis and visualization.

### Platform Overview

HBIcloud offers a suite of 94 tools covering various omics disciplines (**Figure 1**). For genomics, it includes tools for sequence alignment^12^, variant calling^13^, genome assembly, and annotation^14, 15^. In the realm of transcriptomics, it provides tools for differential gene expression analysis, transcript assembly, and functional annotation. Proteomics tools facilitate protein identification, quantification, and functional analysis, while metabolomics tools aid in metabolite identification, quantification, and pathway analysis^16, 17^. For phenomics, HBIcloud offers tools for phenotype data analysis and visualization. Furthermore, the platform includes tools for multi-omics integration, such as clustering, dimensionality reduction, and network analysis^18-20^. Additionally, a collection of mini tools for specific tasks enhances the flexibility of the platform.

HBIcloud is designed with the user in mind, offering features that ensure a seamless experience. Standardized workflows for common analyses reduce the learning curve for new users, while extensive parameter options allow advanced users to fine-tune their analyses. The platform’s graphical user interface eliminates the need for coding skills, making it accessible to researchers from diverse backgrounds. Comprehensive tutorials and a detailed FAQ section assist users at every step, ensuring they can effectively utilize the platform’s capabilities. Moreover, HBIcloud supports various output formats, including publication-quality figures with various formats, and is compatible with multiple input file types, facilitating data import from various sources. The platform also offers instant access to all tools without the need for user registration, promoting ease of use and accessibility.

### Applications and Impact

HBIcloud supports a wide range of applications in multi-omics research. By providing tools for the normalization and harmonization of diverse datasets, it ensures data comparability and reproducibility. Its integrative analysis capabilities enable the combination of different omics layers to uncover complex biological relationships and regulatory mechanisms^21-23^. The platform’s powerful visualization tools present data in an interpretable and publication-ready format, aiding researchers in effectively communicating their findings.

In multi-omics research, the suite of tools available on the HBIcloud platform plays a pivotal role in deciphering the complexity of biological systems, integrating multidimensional data, analyzing statistical relationships, and streamlining analytical workflows. Within the genomics toolkit (**Figure 2**), tools such as Hi-C, Chr. circle plot, and GuideRNA design are particularly crucial for understanding the relationship between chromosomal structure and gene function. The Hi-C technology captures the three-dimensional conformation of chromosomes across the entire genome, revealing interactions within genomic domains and providing spatial insights into gene regulation. The Chr. circle plot tool visualizes chromosomal conformation data, showcasing the three-dimensional structure of chromatin, thereby aiding scientists in identifying functional chromatin loops and their roles in gene expression. The GuideRNA design tool offers precise targeting sequence design for CRISPR-Cas9 gene editing, supporting both genome editing and functional validation.

**Figure 2.**
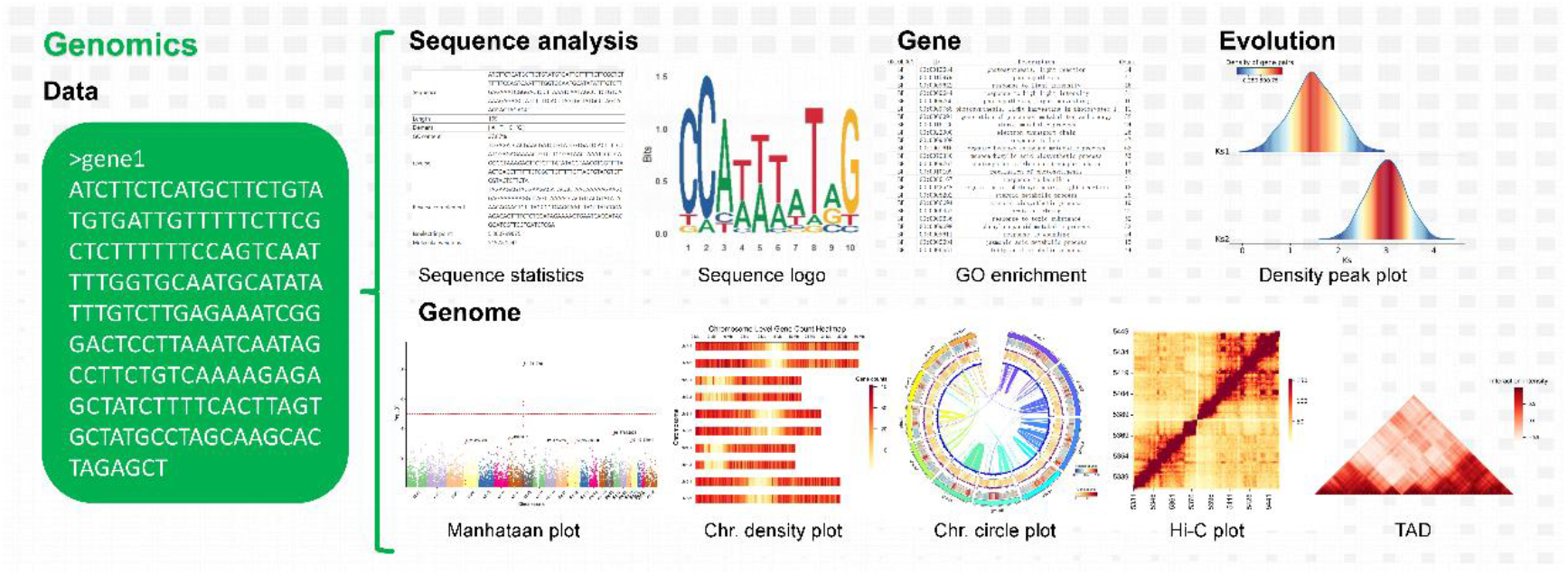
Representative genomic tools in HBIcloud. These include tools for sequence analysis, genomic analysis, genetic analysis, and evolutionary analysis.

In transcriptomics research, tools such as various heatmaps and WGCNA-related plotting analyses have significantly advanced our understanding of gene expression patterns and their functional associations (**Figure 3**). HBIcloud offers a diverse range of heatmaps, including circular, contour, 3D, and dot heatmaps, enriched with extensive visualization capabilities. Most heatmaps on the platform come with clustering options, featuring seven clustering methods and twenty-two distance metrics. These heatmap tools use intuitive color gradients to depict variations in gene expression, enabling researchers to quickly identify shifts in expression patterns and their associations with biological traits. The Module-Trait Analysis tool within WGCNA is particularly crucial, as it uncovers modular features of gene regulation by linking gene expression modules with phenotypic traits. This provides a powerful means to systematically understand the functional mechanisms at the transcriptomic level.

**Figure 3.**
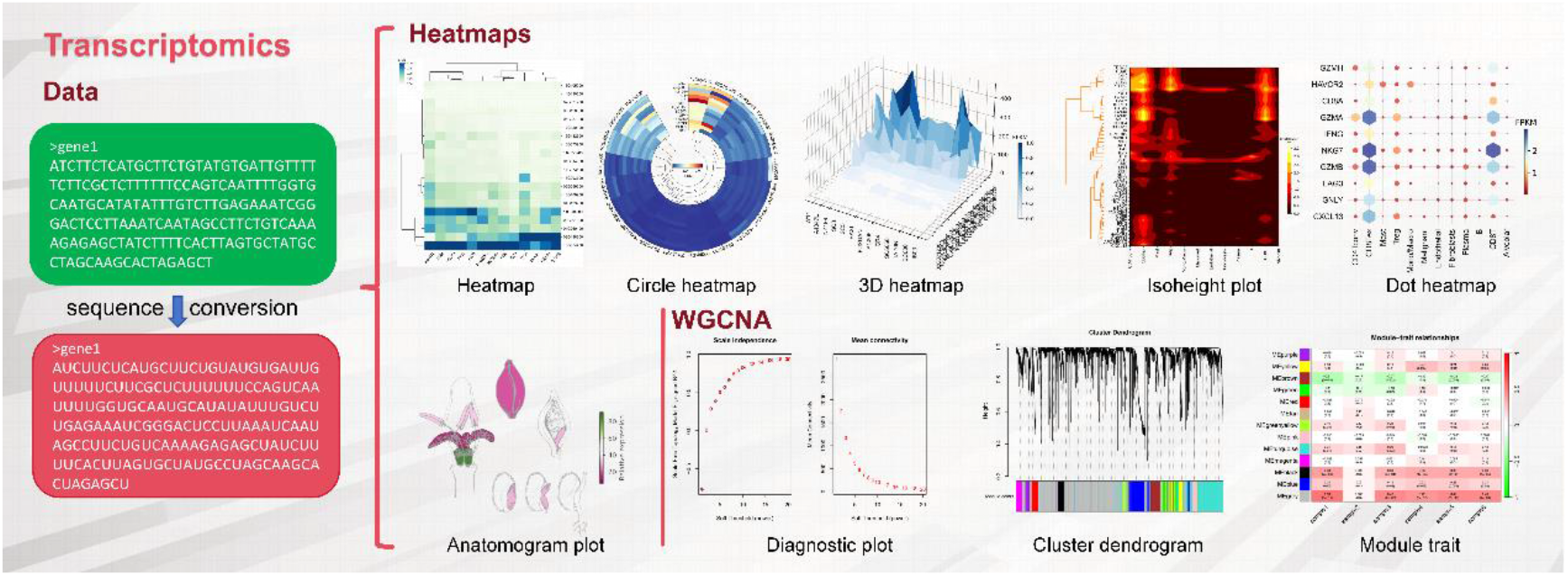
Representative transcriptomic tools in HBIcloud. These include tools for sequence analysis, expressional analysis, and correlation analysis.

HBIcloud offers crucial proteomics tools such as protein analysis, protein domain analysis, and 3D protein structure visualization, which are essential for elucidating protein functions and interactions (**Figure 4**). The Protein Analysis tool provides key metrics such as amino acid composition, molecular weight, aromaticity, instability index, isoelectric point, and molar extinction coefficient, aiding in the investigation of protein biological functions and interactions. The protein domain analysis tool identifies functional domains within proteins, revealing the functional modules of proteins and their roles in biological processes. The 3D protein structure prediction tool constructs and visualizes the three-dimensional conformation of proteins, enabling researchers to comprehend the spatial structure of proteins and their interactions with ligands or other molecules. This facilitates a deeper exploration of protein biological functions and the identification of potential drug targets.

**Figure 4.**
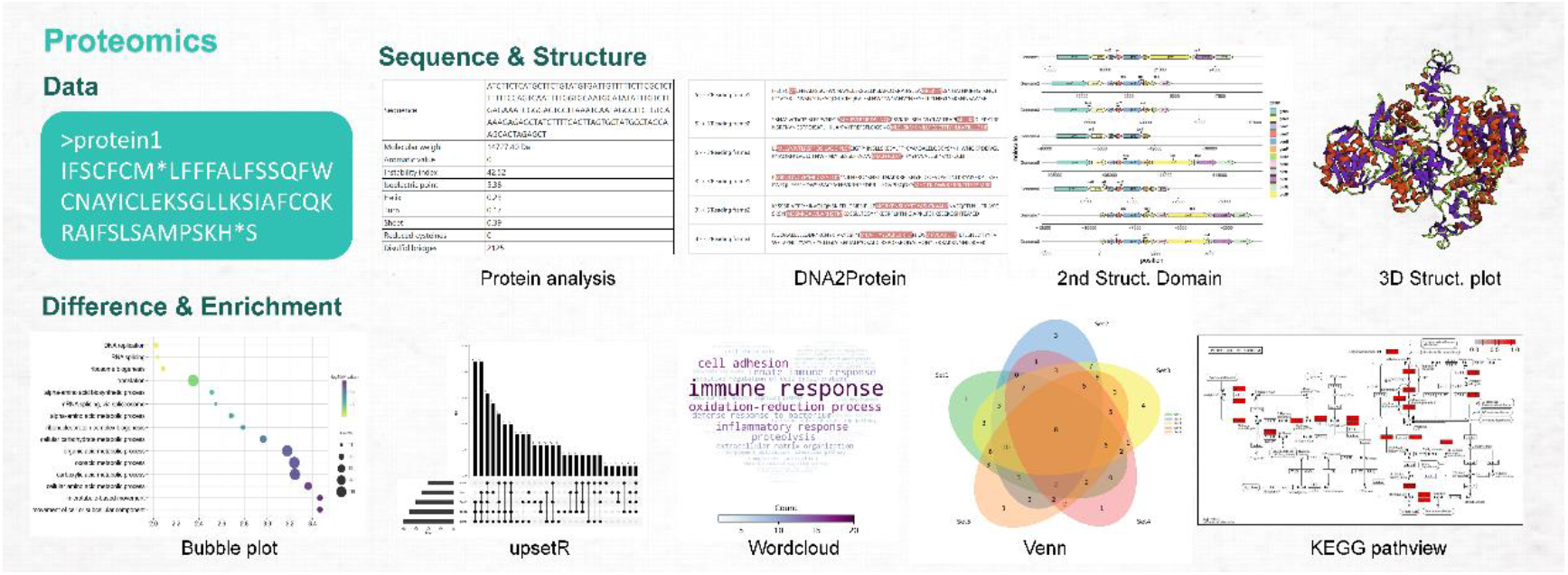
Representative proteomic tools in HBIcloud. These include tools for sequence analysis, annotation, difference and enrichment analysis.

In metabolomics, key tools such as network diagrams, volcano plots, and Corrplot play a significant role in analyzing metabolic pathways and the relationships between metabolites (**Figure 5**). The network diagram tool visualizes the complex interactions within metabolic pathways, revealing critical nodes and key pathways in the metabolic network, thereby providing a comprehensive perspective on metabolic regulation. Volcano plots highlight differential expression and statistical significance of metabolites, aiding in the identification of key metabolites with potential biomarker roles. Corrplot, by displaying the correlation matrix among metabolites, helps uncover synergistic effects and interactions within the metabolic network, offering a powerful tool for the in-depth exploration of metabolomics data.

**Figure 5.**
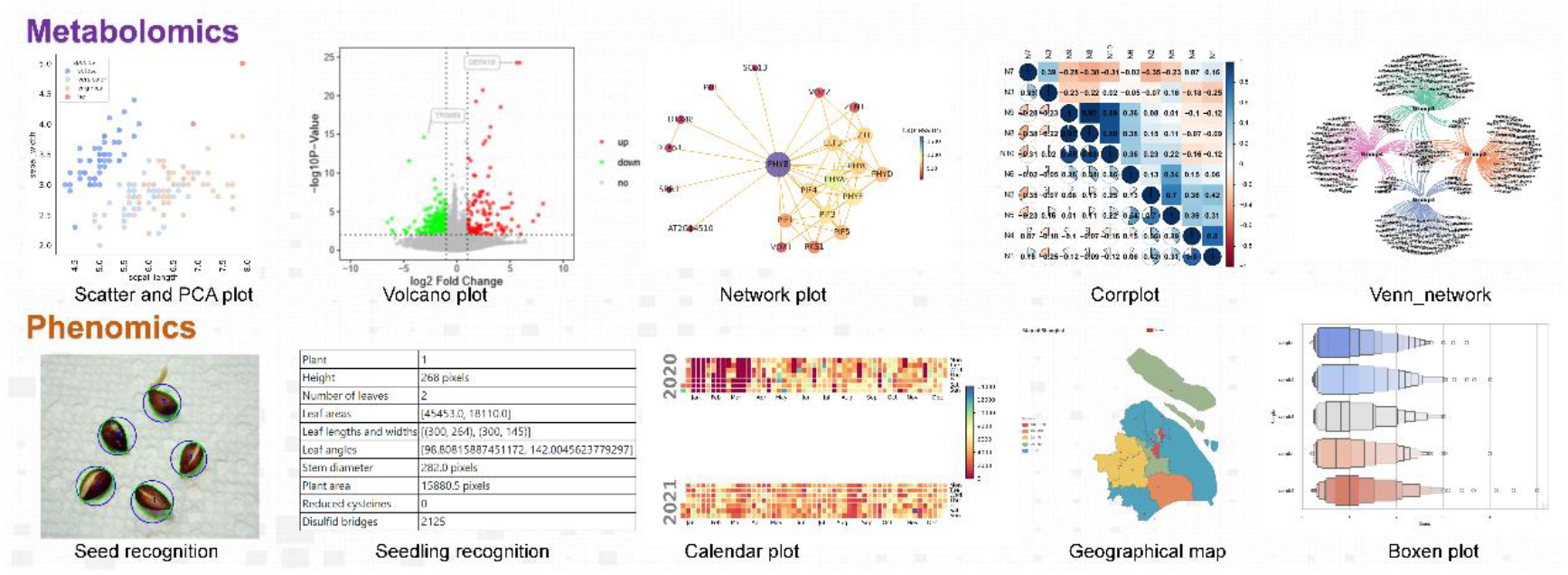
Representative metabolomic and phenomic tools in HBIcloud. These include tools for correlation analysis, picture recognition, and data visualization.

In phenomics, distinctive tools such as Seed Recognition and Seedling Recognition offer valuable insights (**Figure 6**). The Seed Recognition tool efficiently analyzes seed images, revealing phenotypic diversity and its relationship to environmental adaptability. The Seedling Recognition tool accurately identifies the growth stages of developing seedlings. The Geographical Map, a type of heatmap, visualizes data across different geographic locations, helping to understand the distribution and quantity of various plant varieties in different regions.

In multi-omics data integration, tools like Z-score, ComBat, and iCluster play a central role in the normalization and integrative analysis of diverse omics datasets. Z-score standardizes the data, making comparisons between different omics datasets more intuitive and accurate, effectively addressing the issue of varying data scales. The ComBat tool applies batch effect correction techniques to eliminate batch effects from experiments, ensuring the reliability of multi-omics data integration. iCluster, a Bayesian-based multi-omics integration tool, simultaneously considers multiple omics datasets to uncover hidden correlations between them, providing a more comprehensive biological insight.

In the statistical omics analysis offered by HBIcloud, tools like Boxplot, Beeswarm plot, and Sankey plot provide essential means for data visualization and statistical analysis. Boxplots display the median, interquartile range, and outliers, helping researchers identify data distribution characteristics and deviations. Beeswarm plots present data points in a non-overlapping manner, visually conveying the density distribution and dispersion of data, offering a unique approach for visualizing small sample datasets. Sankey plots illustrate the relationships between variables, helping researchers understand complex interactions and flow relationships, making them widely applicable in pathway analysis and resource allocation studies within omics research.

The mini-software modules in HBIcloud offer convenient auxiliary functions for everyday data management and analysis, such as the QR code generator and MD5 checksum tools. The QR code generator allows researchers to quickly convert data or links into QR codes, facilitating fast information sharing and storage, thereby enhancing the efficiency of communication. The MD5 checksum tool verifies file integrity by generating a unique MD5 code, ensuring the accuracy of data during transmission and storage, and protecting against tampering or corruption, providing a fundamental layer of security for data management.

These tools play a unique role in their respective fields, driving the advancement of multi-omics research through diverse omics visualization and analysis, multidimensional data integration, and statistical analysis. They not only enhance data processing capabilities but also provide richer analytical dimensions and technical support for multi-omics research, fostering a more comprehensive understanding of biological systems.

The combination of comprehensive tools, user-friendly features, and robust documentation makes HBIcloud an invaluable resource for researchers. Its ability to facilitate high-quality, reproducible multi-omics analyses positions it as a highly citable platform in the scientific community.

### Future Prospects

Looking ahead, HBIcloud aims to continuously expand its toolset by incorporating emerging technologies and methods. This includes tools for single-cell omics^24^, spatial omics^25^, and advanced machine learning algorithms^26^ for integrative analysis. By staying at the forefront of technological advancements, HBIcloud ensures it meets the evolving needs of the research community.

Enhancing the user experience is another priority for HBIcloud’s future development. Efforts will focus on improving the interface design to make it more intuitive and responsive. Incorporating user feedback will guide regular updates, ensuring the platform addresses the needs and preferences of its users. The development of automated workflows will further streamline complex analyses, reducing manual intervention and enhancing efficiency.

Collaborative features are also planned for future releases. These features will allow researchers to share datasets, workflows, and results within the platform, fostering collaboration and knowledge sharing within the scientific community. This collaborative approach will help build a stronger, more connected research community, driving collective progress in multi-omics research.

**Figure X.**
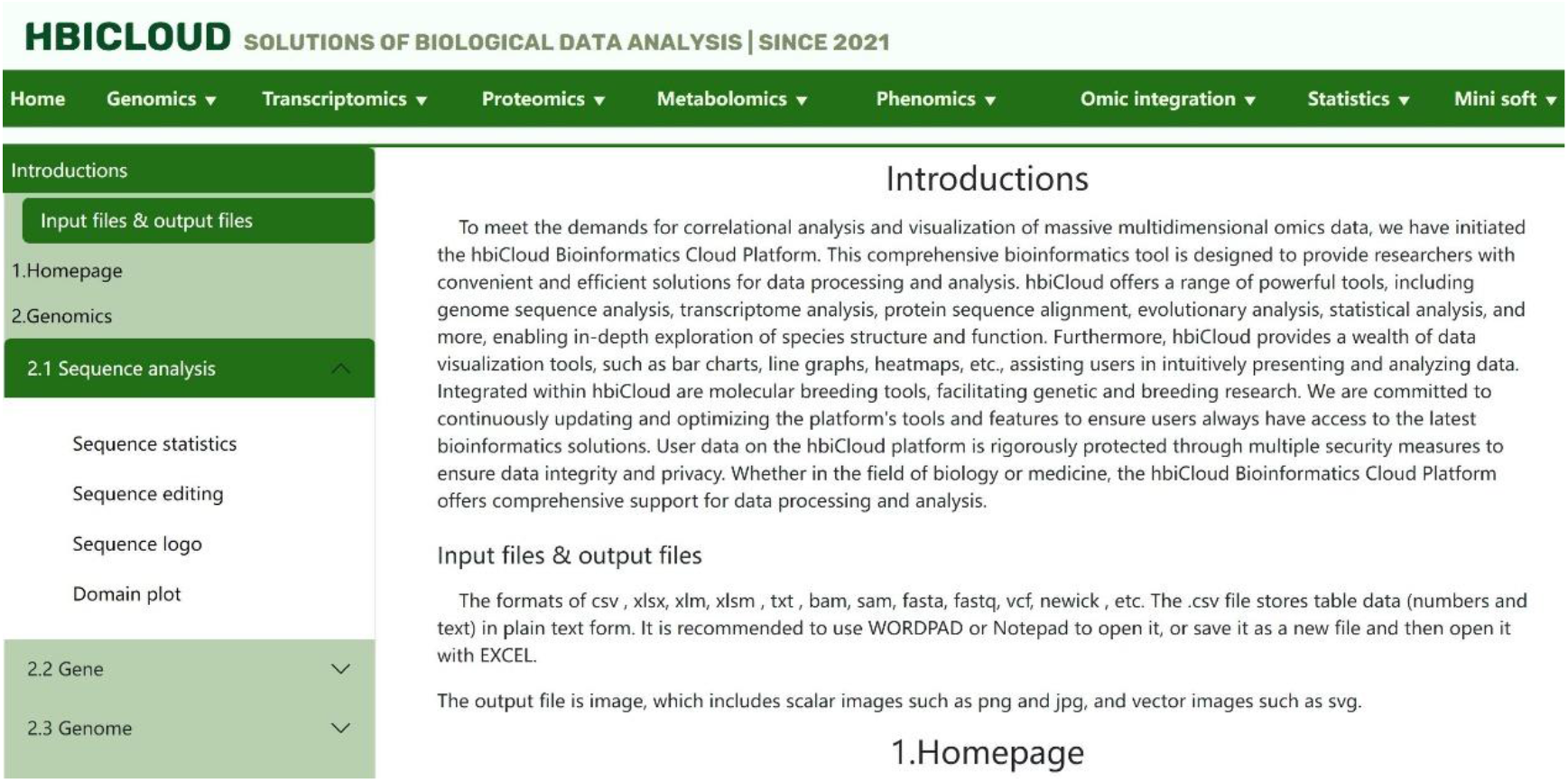
HBIcloud offers comprehensive online tutorial for each tool.

To support the research community further, HBIcloud plans to offer online courses, webinars, and workshops on multi-omics data analysis using the platform (Figure X.). These educational initiatives will promote the adoption and effective use of HBIcloud, ensuring researchers can fully leverage its capabilities for their studies.

## Conclusion

HBIcloud represents a significant advancement in the field of multi-omics data analysis, offering researchers a comprehensive, accessible, and powerful toolkit. The genomics tools provided by HBIcloud facilitate a profound understanding of the complex relationships between chromosomal structure and function, enhancing our ability to systematically decipher gene regulatory networks. By visualizing gene expression patterns and their associations with phenotypic traits, the transcriptomics tools reveal the dynamic networks governing gene regulation, offering insights into the temporal and spatial aspects of gene activity.

In proteomics, HBIcloud’s tools provide deep insights into protein structure and function, advancing our understanding of molecular interactions and applications. These tools elucidate the intricate details of protein conformation and its implications for biological processes, paving the way for novel discoveries and therapeutic strategies. The metabolomics tools offer a multi-dimensional perspective on metabolic pathways, highlighting critical nodes and synergistic effects within metabolic networks. This comprehensive analysis supports systemic understanding of metabolic regulation and its broader implications.

Phenomics tools integrate spatial and morphological data to dissect the genetic basis of plant phenotypic traits, enhancing our knowledge of plant biology and breeding. The Omic Integration tools standardize and integrate multi-omics data, delivering holistic insights into biological big data by bridging disparate datasets and revealing overarching patterns. Statistical omics tools, through diverse visualization and analytical methods, enhance the interpretability of data, making complex information more accessible and actionable.

Additionally, the Mini software tools streamline data sharing and verification processes, boosting research efficiency and ensuring data security. The comprehensive functionality, user-friendliness, and ongoing commitment to improvement position HBIcloud as a leading platform in the multi-omics research domain. By enabling high-quality integrated analyses, HBIcloud holds the potential for significant citation impact and substantial contributions to scientific discovery, marking it as a pivotal resource in advancing our understanding of biological systems.

## Author Contributions

Fei Chen conceived the study. Fei Chen and Wenquan Wang supervised the study. Yunqing Luo implemented the computational pipeline. Shuang He and Wei Dong performed the data analysis and visualization. Fei Chen wrote the manuscript. All authors reviewed and approved the manuscript.

## Acknowledgement

This work was supported by the Hainan Province Science and Technology Special Fund (ZDYF2023XDNY050), Hainan Provincial Natural Science Foundation of China (324RC452), and the Project of National Key Laboratory for Tropical Crop Breeding (NO. NKLTCB202337).

